# Data Fusion for Integrative Species Identification Using Deep Learning

**DOI:** 10.1101/2025.01.22.634270

**Authors:** Lara M Kösters, Kevin Karbstein, Martin Hofmann, Ladislav Hodac, Patrick Mäder, Jana Wäldchen

**Affiliations:** Biogeochemical Integration, Max-Planck-Institute for Biogeochemistry, Jena, 07745, Germany; Data-intensive Systems and Visualization Group dAI.SY), Technical University Ilmenau, Ilmenau, 98693, Germany; German Centre for Integrative Biodiversity Research iDiv) Halle- Jena-Leipzig, 04103, Leipzig, Germany; Faculty of Biological Sciences, Friedrich Schiller University Jena, Jena, 07745, Germany

**Keywords:** data fusion, species identification, deep learning, DNA, images, integrative taxonomy, species confusion

## Abstract

DNA analyses have revolutionized species identification and taxonomic work. Yet, persistent challenges arise from little differentiation among species and considerable variation within species, particularly among closely related groups. While images are commonly used as an alternative modality for automated identification tasks, their usability is limited by the same concerns. An integrative strategy, fusing molecular and image data through machine learning, holds significant promise for fine-grained species identification. However, a systematic overview and rigorous statistical testing concerning molecular and image preprocessing and fusion techniques, including practical advice for biologists, are missing so far. We introduce a machine learning scheme that integrates both molecular and image data for species identification. Initially, we systematically assess and compare three different DNA arrangements (aligned, unaligned, SNP-reduced) and two encoding methods (fractional, ordinal). Later, artificial neural networks are used to extract visual and molecular features, and we propose strategies for fusing this information. Specifically, we investigate three strategies: I) fusing directly after feature extraction, II) fusing features that passed through a fully connected layer after feature extraction, and III) fusing the output scores of both unimodal models. We systematically and statistically evaluate these strategies for four eukaryotic datasets, including two plant (Asteraceae, Poaceae) and two animal families (Lycaenidae, Coccinellidae) using Leave-One-Out Cross-Validation (LOOCV). In addition, we developed an approach to understand molecular-and image-specific identification failure. Aligned sequences with nucleotides encoded as decimal number vectors achieved the highest identification accuracy among DNA data preprocessing techniques in all four datasets. Fusing molecular and visual features directly after feature extraction yielded the best results for three out of four datasets (52-99%). Overall, combining DNA with image data significantly increased accuracy in three out of four datasets, with plant datasets showing the most substantial improvement (Asteraceae: +19%, Poaceae: +13.6%). Even for Lycaenidae with high identification accuracy based on molecular data (>96%), a statistically significant improvement (+2.1%) was observed. Detailed analysis of confusion rates between and within genera shows that DNA alone tends to identify the genus correctly, but often fails to recognize the species. The failure to resolve species is alleviated by including image data in the training. This increase in resolution hints at. a hierarchical role of modalities in which molecular data coarsely groups the specimens to then be guided towards a more fine-grained identification by the connected image. We systematically showed and explained, for the first time, that optimizing the preprocessing and integration of molecular and image data otters significant benefits, particularly for genetically similar and morphologically indistinguishable species, enhancing species identification by reducing modality-specific failure rates and information gaps. Our results can inform integration efforts for various organism groups, improving automated identification across a wide range of eukaryotic species.

---

DNA has established itself as a widely used data source for automated species identification e orts in both ecology and evolutionary research, helping to explore evolutionary relationships and genetic diversity (Stuessy, 2009; Karbstein et al., 2024). Traditionally, DNA-based species identification methods rely on short (approx. 1000 bp) genetic markers known as barcodes that can be queried against large databases for species identification (Hebert et al., 2003a; Ratnasingham and Hebert, 2007). Examples include the NCBI nucleotide or the Barcode of Life initiative (BOLD) databases (Dietz et al., 2023; Wiechers et al., 2023). Barcodes and the use of metabarcoding facilitate the assessment of, e.g., environmental samples from aerobiological surveys to discern plant species through pollen (Leontidou et al., 2021), or analyse biodiversity hotspots (Lahaye et al., 2008; Bessey et al., 2020). Nevertheless, identification based on genetics can provide suboptimal results in cases where sequences are too few or too short due to sampling or sequencing issues, or where genetic regions are less variable due to strong pressures for natural selection (Braukmann et al., 2017; Meiklejohn et al., 2019). Multi-locus analyses have been repeatedly proposed to circumvent apparent drawbacks of single genetic marker-based identification, ranging from two to hundreds of loci (Krawczyk et al., 2014; Dietz et al., 2023). However, multi-locus compared to single-locus analyses increase lab work and sequencing costs, are computationally more expensive, or are sometimes difficult to interpret in case of gene tree conflicts (Karbstein et al., 2022; Dietz et al., 2023). In recent years, machine learning (ML) and, in particular, deep learning (DL) approaches have gained recognition in automatizing DNA-based tasks such as the identification or delimitation of species (Zhang et al., 2008; Derkarabetian et al., 2019), DNA basecalling (Boza et al., 2017), genome assembly polishing (Huang et al., 2021), and phylogenetic tree building (Bhattacharjee and Bayzid, 2020).

Besides genetics, taxonomic research still largely relies on the study of morphological characteristics. ML and, in particular, DL as a branch of ML have become especially popular in the identification of species based on images (Buschbacher et al., 2020; Mäder et al., 2021; van Klink et al., 2022; Green et al., 2023). This advancement can be attributed to DL’s ability to efficiently learn to recognize discriminatory, e.g., visual patterns, which in turn enables the algorithm to evaluate the often extensive, feature-rich biological datasets like diverse images of species (Wäldchen and Mäder, 2018).

Additionally, DL algorithms are becoming increasingly prevalent in image-based species identification due to the availability of seminal network architectures such as ResNet (He et al., 2016), which o er scientists a solid foundation for a myriad of derived applications, especially in the automatic identification of different species groups (Norouzzadeh et al., 2018; Mäder et al., 2021; Høye et al., 2021).

Image-based identification can either utilize *in situ*, i.e., field recorded (Terry et al., 2020; Rzanny et al., 2022), specimens or preserved specimens from collections (Carranza-Rojas et al., 2017; Marques et al., 2018; Hodač et al., 2023). The availability of *in situ* images is rapidly increasing, particularly due to citizen science initiatives (Boho et al., 2020; Mesaglio et al., 2023). Preserved specimens, represented by millions of samples in natural history collections (Bebber et al., 2010; Scott and Livermore, 2021), have become more relevant through increasing e orts to automate digitization (Blagoderov et al., 2012; Tegelberg et al., 2014). Compared to genetics-based approaches, images from either *in situ* or collection material provide a fast and low-cost means to species identification that can reliably discriminate between species with a characteristic morphology (e.g. Mäder et al., 2021; Shirai et al., 2022). Nevertheless, image-based species identification comes with its own set of weaknesses, mainly introduced by cryptic species, high phenotypic plasticity, and multiple origins of the same morphotype. These factors can also cause discrepancies between systematics inferred from genetics and morphology, leading to revisions of former morphospecies (e.g. Karbstein et al., 2020; Marcussen et al., 2022). Highly variable image qualities and ways of recording further complicate, identification and can lead to poor identification accuracy, especially for species that are intrinsically hard to distinguish (Wäldchen et al., 2018; Barbedo and Castro, 2019; Chiu et al., 2020).

To overcome the limitations of species identification using a single data point per specimen, multiple data points either of the same modality or of different modalities can be Combined (e.g. Terry et al., 2020). Fusion of data can leverage machine learning algorithms, which are adept at efficiently integrating disparate information (Karbstein et al, 2024). Across disciplines, some attempts have been made to incorporate multiple inputs into ML training. For example, the fusion of different image perspectives yields enhanced results in terms of species identification accuracy (Marques et al., 2018; Rzanny et al., 2022). On the other hand, integrative taxonomy seeks to overcome the limitations outlined above by incorporating various types of data, thereby reducing modality-specific (i.e., data type-specific) failure rates and information gaps (Dayrat, 2005; Schlick-Steiner et al., 2010; Karbstein et al., 2024). While its utility has been limited by the large amount of data required since traditional procedures rely on extensive pipelines that often include manual labour and do not scale well, the application of machine learning methods o ers a solution. For instance, recent developments involve fusing with supplementary metadata such as location or date (Terry et al., 2020). Nevertheless, particularly the integration of DNA and image data emerges as a promising route to high-accuracy species identification. So far, deep learning-assisted fusion of DNA and images has primarily been applied in biomedical research (Stahlschmidt et al., 2022). To date, few studies have used genetic and image input for species identification. Yang et al. (2022) have developed a new convolution-based architecture for integrative species identification (MMNet) using barcodes and images. They found that MMNet outperformed existing methods on 10 distinct datasets comprising both animal and plant groups while achieving very high accuracies with up to 100% identification success. Badirli et al. (2023) used a hierarchical Bayesian model to integrate DNA and image features of four insect orders derived from unimodal convolutional neural networks (CNN). They found that multimodal species identification performed better than unimodal ML methods but was surpassed by traditional distance-based identification on DNA data. In addition, notable e orts in species delimitation include the successful unsupervised training of a SuperSom, i.e., an ANN producing a multi-layer grid, where each layer represents an input type (Alexander Pyron, 2023). Another example is the use of a Bayesian approach that can incorporate multiple loci and quantitative traits to suggest alternatives for provided species labels (Solís-Lemus et al., 2015). Guillot et al. (2012) employ a statistical approach with the goal of building homogeneous clusters without needing prior knowledge and using spatial, phenotypic, and genetic information. Our exploration of the fundamental question of how to preprocess genetic data for use with DL models and how to fuse genetic data with other types of input extends the aforementioned studies.

Generally, DNA has to be specifically preprocessed in order to work with ML methods. Multiple options of DNA arrangement and numerical encoding are imaginable. Researchers can either choose to input sequences in their raw state, align them, or additionally reduce them to single nucleotide polymorphisms (SNPs). DNA must also be transformed into a numerical representation by turning each base into a vector of a chosen length *n*, by assigning a specific numerical value to each base, or by learning an informative representation. Such a representation can be learned using, for instance, similarity learning approaches or, alternatively, a transforms architecture. Notable examples of DNA-focused transformers are DNABERT (Ji et al., 2021) and DNAGPT (Zhang et al., 2023). Transformers are powerful encoder-decoder structures that are applied to text-like input. For comparability between the two modalities images and DNA, we have opted to use CNNs instead. Additionally, there is a diverse range of deep learning fusion methods (e.g. Seeland and Mäder, 2021). These methods encompass techniques such as feature fusion, where features extracted from different modalities after any but the last layer are combined to provide a more comprehensive representation of the data. Moreover, score fusion aggregates predictions from multiple models or modalities to make a final decision (Seeland and Mäder, 2021). To date, there has not been a systematic and statistical comparison of DNA preprocessing and multimodal fusion approaches combining genetic and image input applied across eukaryotes. Nonetheless, the choice for a preprocessing and fusion method can enhance or limit both the achievable identification accuracy and the efficiency of the model. For instance, reducing the DNA to SNPs can decrease noise (e.g., missing data/gaps) and greatly accelerate training times when working with very long sequences. Determining preprocessing and fusion methods that yield consistently robust results across eukaryotic groups is, therefore, critical for species identification e orts using ML. In addition, a comparison to the baseline, i.e., the accuracy achieved by relying on a single datatype, is of importance as fusion always involves additional e ort and, thus, should only be considered when accompanied by a considerable increase in identification accuracy. In this study, we use the genetic markers *COI* for two animal and *rbcLa* for two plant families in combination with image-based morphological data to systematically investigate preprocessing and fusion methods. In addition, we want to provide future studies on integrative systematics with a baseline to guide them during sample collection and the process of choosing an appropriate model architecture. Specifically, we explore four key targets: 1) the determination of the DNA arrangement and 2) encoding options that yield the most accurate species identification across diverse datasets, 3) investigation of the impact that data fusion has on species identification success, 4) assessment of the effectiveness of the proposed fusion strategies. Finally, we explore the mechanisms underlying accuracy improvements resulting from the combination of genetic and image data as a contribution to explainable Al.

## Material and Methods

### Dataset collection and Itering

We assembled four datasets, each comprising DNA and image data: two plant datasets and two animal datasets (Table 1). Each dataset focuses on a specific family, with varying numbers of genetic distances between and within genera and species to present various levels of complexity with regard to differentiation. We gathered data from the Asteraceae and the Poaceae family for the plant datasets. In the Poaceae dataset, we utilized images captured in natural habitats, while for the Asteraceae dataset, we relied on digitized herbarium material. This approach enabled us to examine the viability of our method for *in situ* plant images as well as for digitized preserved specimens. The animal datasets consist of the Coccinellidae and the Lycaenidae family. Due to data availability constraints, we could only utilize images from preserved specimens, i.e., from collections, for both datasets. For the same reason, we chose to set the minimum number of records per species to four for all datasets, where each record is composed of a molecular and an image sample. The low number of records reflects the reality many researchers face in their sampling e orts (e.g., compare to Zarrei et al., 2015; Karbstein et al., 2020; Yang et al., 2022). Although other papers such as (Badirli et al., 2023) have already collected datasets comprising molecular and image data, we opted to gather our own. This decision was motivated by our need for multiple datasets from both plants and animals, the intention to include *in situ* and preserved material, and the nature of our research questions. Since we were not interested in the total success rates of our models, but rather the relative gains and losses between single- and multimodal trainings, we did not include any comparisons to architectures proposed by other research papers. We have used the genetic markers *COI-5P* for animals and *rbcLa* for plants. Although genome-level data would have been an alternative to using genetic markers, they entail multiple challenges, First, DNA barcoding has been a widely used method for species identification due to its relative efficiency in terms of time, cost, and resources, which lead to extensive data collections and the establishment of dedicated databases, such as BOLD (Ratnasingham and Hebert, 2007).

**Table 1.** Dataset overview. Datasets vary in the number of genera, species, and samples within the respective family. We relied on either *in situ* images or pictures from preserved specimens (collections). Information on the number of samples with related image and DNA data (combined) is listed alongside total sample size. We used *COI 5P* for both animal and *rbcLa* for the plant datasets. Mean gene lengths as well as standard deviations are listed as well. M=median. =mean

| Family | No. of samples (combined) | Samples per species (M) | No. of species | Species per genus (M) | No. of genera | Barcode | Gene length ( ) | No. of SNPs | Image type |
| --- | --- | --- | --- | --- | --- | --- | --- | --- | --- |
| Asteraceae | 970 (447) | 5 | 146 | 1 | 45 | *rbcLa* | 550 ( 13) | 106 | collections |
| Poaceae | 1118 (0) | 7 | 123 | 1 | 54 | *rbcLa* | 555.9 ( 22) | 114 | *in situ* |
| Coccinellidae | 1092 (683) | 9 | 72 | 1 | 33 | *COI 5P* | 618.4 ( 59) | 398 | collections |
| Lycaenidae | 5520 (2559) | 8 | 259 | 1 | 98 | *COI 5P* | 645.3 ( 45) | 482 | collections |

When selecting genetic markers, we considered both data availability and resource efficiency, as well as how these choices would enhance the study’s value for future research. Second, additional formatting questions arise when working with genome-level data and CNNs. Genome-level data is much more high-dimensional than the purposefully short genetic markers and, thus, is more suited as input to alternative architectures and attention mechanisms (e.g., transformer) that can digest genome-level data more effectively. However, in this study, we actively decided on CNNs for their use in both image-based and DNA-based research. Implementing a CNN for both modalities facilitated an easy and valid comparison between models.

#### Barcodes

We sourced all genetic data from publicly available repositories, namely BOLD (Barcode of Life Data Systems) and GenBank via their APIs. Our queries primarily relied on the family name. Since BOLD stores image and molecular data, we searched for combined records, i.e., ‘specimen’ and ‘sequence’. GenBank does not o er image data. Thus, within our GenBank queries, we focused on ‘genomic DNA/RNA’ (property) and ‘gene or RNA’ (feature key) data. Inspired by Paris et al. (2017) and Karbstein et al. (2020, 2021), we decided to assess and use sequence clustering and alignment features for filtering and to ensure a reasonable similarity between same-locus sequences. First, we ensured duplicate removal of GenBank records that were already sourced from BOLD by checking the GenBank accessions. Within each dataset, we then selected the five most abundant genetic markers. All associated sequences underwent clustering, with sequence similarity thresholds ranging from 0.5 (low similarity) to 0.99 (high similarity) in 0.01 increments using VSEARCH v2.22.1 (Rognes et al., 2016). To be able to keep as many samples as possible, we conducted all further calculations on the largest cluster per threshold. We determined the number of species and samples in this cluster as well as generated an alignment with MAFFT v7.490 (Katoh and Standley, 2013) to compute the SNP and gap counts relative to the aligned sequence length using SNP-sites v2.5.1 (Page et al., 2016) and VCFtools vO.1.16 (Danecek et al., 2011). For each category (i.e., SNP and gap), we then calculated the differences in values between the score of one threshold *x* and the optimal score *y*. The optima represented the highest SNP and the lowest gap count across all thresholds. To combine both aspects - SNP and gap counts - we calculated a weighted mean, with double the emphasis on reducing the gap score. Finally, we chose the threshold with the lowest combined divergence. Choosing a threshold based on a high SNP and a low gap score allowed us to maximize the information content while ensuring that our sequence clustering was correct.

Based on the chosen thresholds for each marker, we then decided on the genetic marker for the respective dataset by considering the number of species retained in the respective final datasets based on the chosen cluster and the average sample size per species. Naturally, a high number of species leads to a more complex learning task while the number of samples strongly impacts how well the model is able to distinguish between classes, i.e. species (e.g. Durden et al., 2021). In detail, all values were sorted within their respective property group, i.e., species number, and sample size. Based on these sorted lists, we assigned numeric indices to each marker (e.g. first position in *species number* and second in *sample size)*. We picked the genetic marker with the lowest averaged index, i.e., the marker presenting the best possible balance between information content and inherent task complexity introduced by many possible species. Our filtering resulted in the choice of *COI-5P* for Coccinellidae and Lycaenidae, and *rbcLa* for Asteraceae and Poaceae with identity thresholds set to 0.78, 0.88, 0.97, and 0.96 respectively. For clustering, we set the maximum sequence length to 660 bp for *COI-5P* (Hebert et al., 2003b) and 670 bp for *rbcLa* (Dong et al., 2014) respectively, while the minimum was set to 40% of the maximum length. All sequences outside that range were discarded.

We observed duplicate sequences in all four datasets, As this is a known obstacle to barcoding, especially in plants (Fazekas et al., 2009), we decided to regard duplicates as naturally occurring identical sequences to more accurately represent the real world instead of removing them from the dataset.

#### Images

In addition to images obtained from BOLD, we collected further images using the GBIF API. Similarly to the search for genetic and combined samples in BOLD, our GBIF search was also based on the dataset family name. For the herbarium material search in GBIF, we specified the basis of the record as ‘preserved specimen’ and ‘material sample’. In regards to *in situ* images, since most images deposited in BOLD depict herbarium specimens, we decided to eliminate all BOLD-derived images before querying other databases. We restricted GBIF results to iNaturalist research-grade observations as well as including images from Flora Capture (Boho et al., 2020) to maximize the trustworthiness of observations. Images from ‘collections’ belong to museums and museum-acknowledged private collections that were labeled by experts instead of citizen scientists as is the case for iNaturalist observations.

Duplicate images were automatically removed in both training and validation sets. Regarding our validation sets, we ensured the following criteria in regards to image quality: The image had to be of the correct type (i.e. showing *in situ* or preserved specimen) and the sample needed to be the focus of the image. Images of low quality, i.e. visibly pixelated images or images with no specimen depicted, were manually removed. Besides the removal of duplicate images, we did not apply any filtering on the training sets. Our reasoning for skipping further filtering steps on the training sets was as follows: A prominent goal when leveraging online databases is to cut down on manual labor. Thus, being able to trust that their derived large, heterogeneous datasets are of reasonable quality and can excel in the context of ML is a vital aspect of future ventures into ML-assisted species identification.

### Experimental setup

#### Model architectures and BLAST

Both images and barcodes were trained using the ResNet-50 architecture (He et al., 2016). ResNets, termed for their residual connections, are seminal and widely adopted convolutional neural networks (CNNs) often chosen as a baseline in computer vision projects (e.g. Mathur and Goel, 2021; De Nart et al., 2022). CNNs are also prevalent in DNA-based research (e.g. Yang et al., 2022; Liu et al., 2022). Here, we have used all 49 convolutional layers of the ResNet-50 as the feature extractor for both unimodal model portions. Thus, our feature extractors have 23.5M parameters. For the ordinal encoding, we prepended an embedding layer, composed of a 1×1 convolution with a sliding window of 1 and no padding. The model’s input channel size was set to four to accommodate the size of our fractional encoding vector and to one for the ordinal encoding. The classifiers for our separate models have 1.8M parameters. For comparability, we adjusted the number of parameters for the fused models to be twice the size of the individual models. We have implemented all models using the framework PyTorch vl,13.0+cull7 (Paszke et al., 2019) under Python v3.9.13. We trained on a Tesla V100-SXM2 and a NVIDIA A40 GPU. Species identification was also performed using *blastn* with default parameters (Camacho et al., 2009). To avoid bias from the extensive records in the NCBI GenBank repository, locally created BLAST databases were used with the BLAST+ *makeblastdb* command.

#### Training

To prevent overfitting, we applied early stopping, i.e., stopping on the training epoch that best generalized over the validation set, with a patience of 20 epochs (Prechelt, 1998; Ying, 2019). The maximum number of epochs was set to 500. Following Seeland and Mäder (2021) and derived from standard values, we applied categorical cross-entropy loss with Adam as the optimizer, using an initial learning rate of le-4, and set our mini-batch size to 32. Due to sample size constraints, we merely relied on the validation set for performance evaluation. We validated model performances on two records per species, All samples beyond that belong to the training sets. For images, we followed standard procedures and applied common augmentation techniques (RandomResizedCrop, RandomVerticalFlip, RandomHorizontalFlip). As a side effect of RandomResizedCrop, most of the time, the model does not encounter the relatively small, but potentially informative labels included in the images of preserved specimens. Since we were only interested in determining which fusion approach performed best, label learning also did not impact our conclusion. We optimized our Random Forest classifiers by applying grid search to each of the datasets beforehand (Text SI). Our decision to optimize the Random Forest was based on the much less time-consuming nature of the algorithm and the meager results we attained in a preemptive test compared to the FC-based classifiers.

Our procedure included training on barcodes and images separately at first to obtain a baseline for comparison. In image classification, it is a common procedure to apply fine-tuning to pre-trained networks (e.g. Mathur et al., 2020; Kirbaş and Çifci, 2022). Here, our image model leveraged the ImageNetlK_V2-trained weights (Deng et al., 2009). When training the multimodal model, we reused the weights acquired by the unimodal training for both image and DNA data. We opted to freeze all layers but the last block of the feature extractor during the second training step to concentrate on higher-level feature learning (i.e., complex visual structures).

To test whether any differences in accuracy between the best fusion approaches and the DNA-and image-only models were statistically significant, we implemented a Leave-One-Out Cross-Validation (LOOCV) for all datasets (see Brownlee, 2020, for detailed explanation). LOOCV is defined by running the model once for each sample in the dataset. Every training uses the entire dataset except for one sample. Validation is then performed on the excluded sample. However, due to time constraints, we decided to validate on a subset of four samples per species for LOOCV. For example, we trained 4 (samples) times 146 (species) models for Asteraceae, summing up to 584 models per uni-/multimodal method in total. Each model utilized all but one sample for training and evaluated on the excluded sample. An alternative to using LOOCV would have been, e.g., a K-Fold Cross-Validation (K-Fold CV). With K-Fold CV, the dataset is split into K folds or subsamples, where all but one fold are used for training and one fold is used to evaluate the model’s performance. Consequently, LOOCV is an extreme version of K-Fold CV. While both approaches have their pros and cons, the very limited number of records per species was the main reason we decided on LOOCV. Employing this approach, we also conducted LOOCV for BLAST, creating the databases from all but one sample per dataset and then querying the left-out sample against the database. Only the first listed hit was considered for the accuracy calculation. To further test the stability of our limited LOOCV, we conducted a more thorough LOOCV for one plant (Asteraceae) and one animal (Coccinellidae) dataset that tested on all samples instead of the subset of 4. The results are shown in Figures S1-S6.

#### Metrics and statistics

We evaluated the performance of our models by means of validation accuracy, i.e., the ratio of correctly classified samples to all samples in the dataset. To check for significant differences in non-normally distributed accuracies within all datasets, we employed the non-parametric, binary Cochran’s Q test for datasets characterized by more than two group factors and paired data using the R package RVAideMemoire (Herve, 2023). Then, we performed pairwise group comparisons based on the McNemar test to investigate differences in detail. Using R v4.3.1 (R Core Team, 2023) and the caret package (Kuhn, 2008), we calculated confusion matrices based on our image, barcode, and best multimodal model predictions to provide a basis for subsequent analyses. We then determined confusion rates and grouped them by intergeneric and intrageneric confusions. For the statistics on confusion rates, we applied the non-parametric Kruskal-Wallis test for paired data in combination with pairwise Wilcoxon signed rank tests. In addition, generalized linear models (GLMs) with binomial response were used to check whether confusion rates are related to barcode (gene) length and sample size using a custom R package. Using, in our case, confusion rates (species-level and derived genus-level) as the response variable and different gene length features (min, max, median) and sample size indices as the predictors, a custom package automates model simplification. To avoid disturbance of modeling procedures (Dormann et al., 2013), the main package function removes autocorrelated variables beforehand (r>0.8; i.e., general gene features and sample size indices to those indices in the training set). The corresponding R scripts together with a README describing their explicit functions will be available upon publication.

### Multimodal species identification scheme

The workflow of our proposed multimodal species identification is illustrated in Figure 1. It comprises three main steps: A) preprocessing of the DNA data, B) unimodal species identification that serves as baseline and C) multimodal identification after different fusion approaches of DNA and image data.

**Fig. 1.**
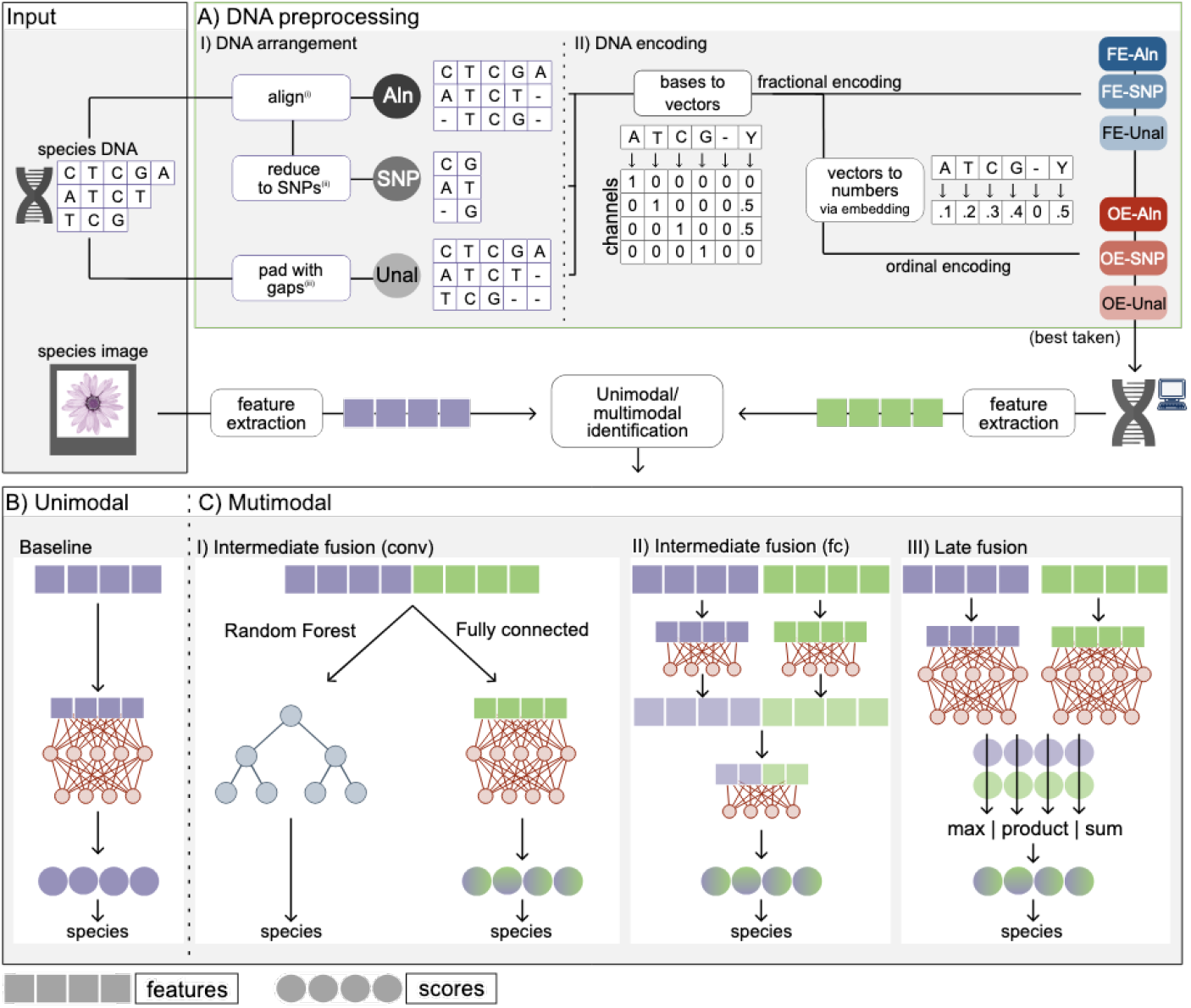
Overview of the multimodal species identification scheme. A) DNA preprocessing route with different I) DNA arrangement and II) DNA encoding methods yielding six variations of the genetic model input in total. B) Unimodal identification with either image or DNA serve as baselines. C) Different fusion approaches for multimodal species identification I) image and DNA fusion after feature extraction (Intermediate fusion (conv)), II) after the first fully connected layer (Intermediate fusion (fc)), and III) after the second fully connected layer (late fusion). SNP = Single Nucleotide Polymorphisms, conv=convolution, fc=fully connected. (Preprocessed) DNA and flower images are from https://pixabay.com (free to use under the Content License).

#### DNA preprocessing

For automated species identification using DNA and deep learning technologies, it is crucial to prepare the genetic data for input (arrangement) and convert it into numerical representations (encoding). We refer to the combination of both as DNA preprocessing. We examined three arrangements for genetic datasets: (i) aligning DNA sequences, (ii) further reducing aligned sequences to SNPs, or (iii) padding the DNA with zeros to the same length instead of aligning. As a result, we obtained three sequence variants referred to as ‘aligned’, ‘aligned-SNP’, and ‘unaligned’ (Fig. 1 A(I)).

The arranged sequences are then encoded numerically before being subjected to deep learning models (Fig. 1 A(II)). One approach is the fractional encoding, which is a variant of the commonly used one-hot encoding method. For one-hot encoded representation, the input sequence is represented by a 4 L matrix, where 4 is the size of the bases vocabulary (A, T, C, and G) and L is the length of the sequence. Each position in the sequence corresponds to a vector of length four, with a single non-zero element representing the bases at that position. Specifically, the bases A, T, C, and G are encoded as four one-hot vectors:: [1,0,0,0], [0,1,0,0], [0,0,1,0], and [0,0,0,1]. Therefore one-hot encoding transforms DNA sequences into binary images with four channels corresponding to A, C, G, and T. The special form of fractional encoding allows for values between 0 and 1. For instance, we encoded T as [0,1,0,0], C as [0,0,1,0], and Y (i.e., T or C) as [0,0.5,0.5,0]. Gaps are encoded as zeros instead of extending the vector to maintain consistency between aligned and unaligned sequences.

The second encoding method is an ordinal encoding, where decimal numbers ranging from 0 to 1 are assigned to each of the bases. Instead of manually selecting them, we let the model learn suitable decimals (see Model architectures for details). The optimal combination of arrangement and encoding methods serves as the input for the subsequent multimodal identification step.

#### Unimodal baselines

To obtain a baseline with which to compare the results of the multimodal approaches, genetic data and images were first trained individually. Here, we extracted the features after the last CNN layer from the image- and DNA-trained models. Subsequently, we passed the features through a classifier consisting of two fully connected layers (see Fig. 1B). Additionally, the traditional method BLAST (Altschul et al., 1990) was applied to assess whether it outperforms ML approaches for DNA-based species identification.

#### Multimodal fusion approaches

Multimodal fusion in our study refers to conducting a combined analysis of molecular and image data to investigate whether the combination yield a more accurate species identification result. Data fusion can be implemented at different stages within the model architecture, Recently, it was demonstrated that approaches fusing multiple image perspectives late in the network typically perform better than those that fuse at an early stage (Seeland and Mäder, 2021). Similarly, in their review on unimodal and multimodal fusion in the biomedical field, Stahlschmidt et al. (2022) noted that early fusion approaches often underperform when dealing with heterogeneous modalities. Consequently, we have combined the separate models within the last two layers of the network.

Specifically, we have fused image and barcode features I) directly after the last convolutional layer (‘intermediate fusion (conv)’), II) after the first of two fully connected (i.e., dense) layers (‘intermediate fusion (fc)’) and III) after the final dense layer (‘late’) responsible for generating the output scores (see Fig. 1C). Here, we expand on the terminology used in Stahlschmidt et al. (2022). Instead of the two dense layers in the intermediate fusion approach, we also employed a Random Forest (RF) using the scikit-learn library to o er an easy and fast-to-train alternative to a fully connected (fc)-based classifier (Pedregosa et al., 2011). Despite not being classified as neural networks, they are highly capable of approximating any function and can learn non-linear relationships (Hastie et al., 2009). In the score-level fusion approach, we examined three methods: sum, product, and max score-level fusion. Therefore, we analyzed a total of six different multimodal identification scenarios per dataset.

## Results

### DNA preprocessing methods

Overall, DNA-based species identification was more accurate in the two animal datasets (i.e., Coccinellidae, Lycaenidae) compared to the plant datasets (Asteraceae, Poaceae; Fig. 2). Within the datasets, we observed significant differences in identification accuracy between the arrangement and encoding methods. Notably, these differences exhibit consistent patterns between all families. In both plant families, all arrangement methods with ordinal encoding proved to be inferior compared to their respective fractional encoding counterparts, with mean accuracies surpassing those of ordinal encoding by 11% for Asteraceae and 13% for Poaceae. Within fractional encoding, padding unaligned sequences resulted in the lowest identification rate (Asteraceae: 44.2%; Poaceae: 72%). In Poaceae, there was no significant difference between aligned and SNP-reduced sequences, whereas in Asteraceae aligned sequences significantly outperformed SNP-reduced sequences by 6%. The animal datasets yielded significantly higher identification accuracy for aligned and SNP-reduced fractional encoded sequences than for the remainder of the arrangement and encoding options (>6% improvement). The three ordinal encoded and the unaligned fractional encoded barcodes provided similar results.

**Fig. 2.**
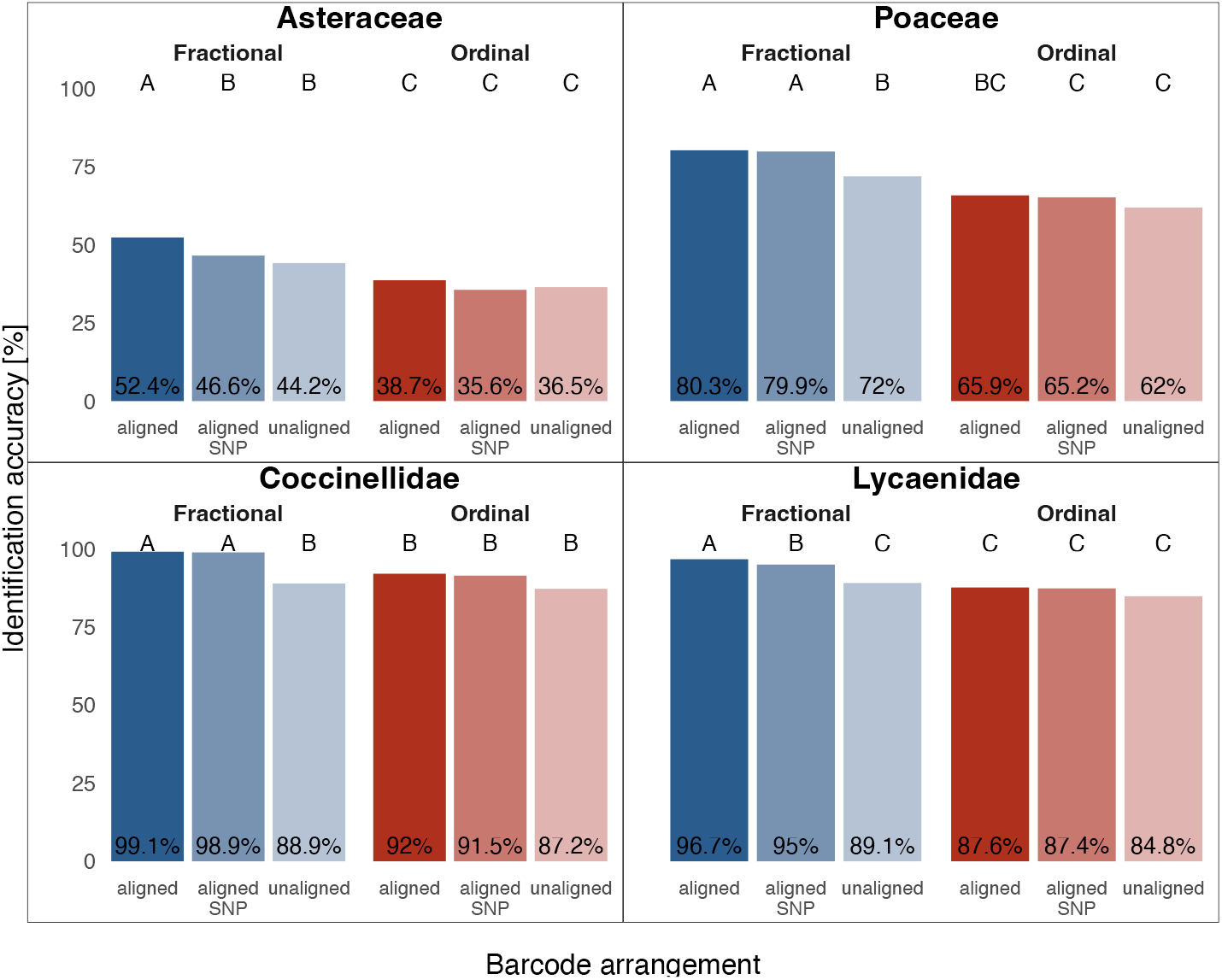
DNA-based species identification accuracy after different arrangement (aligned, aligned-SNP, unaligned) and encoding (fractional, ordinal) methods for genetic data. Identification accuracy describes the percentage of samples within the validation set correctly identified by the model. Letters indicate significant differences in performance (paired Cochran’s Q and pairwise McNemar’s tests, p<0.001). SNP = Single Nucleotide Polymorphisms.

Similarly to Poaceae, there was no significant difference between aligned and aligned-SNP sequences in Coccinellidae. In Lycaenidae, aligned sequences exceeded aligned-SNP sequences in identification success (96.7% compared to 95%).

In both animal and plant datasets, fractional encoding performed better for aligned and aligned-SNP sequences than ordinal encoding (p 0.0001). For unaligned sequences, fractional encoding was significantly better for the two plant groups (p 0.0001) and the Lycaenidae family (p 0.01), whereas in the Coccinellidae family no significant differences could be determined. For fractional encoding, unalignment resulted in a severe dip in identification success compared to alignment (p 0.0001). Furthermore, the magnitude of the differences in accuracy between the preprocessing methods varied between datasets. For example, in Poaceae and Coccinellidae, the biggest margin between two preprocessing methods was between fractional encoded aligned sequences and ordinal encoded unaligned sequences with 18% and 12%, respectively.

An expanded LOOCV that included all samples within the Asteraceae and Coccinellidae datasets confirmed the results found when using the subsets. The only difference observed was a minor change within the ordinal encoded sequences within the Coccinellidae dataset, which shows that the unaligned sequences performed significantly worse than the other two arrangements (Fig. S1).

### Unimodal species identification

In a first step towards fusion and to obtain a baseline for the evaluation of our fusion approaches, we trained on each modality separately. We observed significant differences between models trained on images or DNA data across all datasets (p 0.0001, Figure 3). For the Poaceae. dataset as well as the two animal datasets, the image-based model was inferior to the one trained on DNA data. However, for the Asteraceae dataset, the identification accuracy achieved by the model trained on images significantly exceeded the DNA-only model, with 66.3% compared to 52.4%, respectively. For the Poaceae dataset, DNA-based identification surpassed image-based identification by roughly 20% (80.3% and 60.6%, respectively). Regarding the Coccinellidae, DNA-based identification achieved species identification with 99% accuracy, whereas images achieved sub-optimal results with 81.3%. Similarly, in the Lycaenidae family, DNA data yielded 96.7% identification accuracy, while the image-based model identified 82.4% of samples correctly, In addition to our DL approach, we investigated the performance of BLAST, the traditional method for DNA-based species identification. BLAST’S performance varied substantially across datasets. In the Asteraceae dataset, it misidentified more samples than the ML unimodal image or DNA approaches, with only 33.4% correctly identified. In Poaceae, it performed similarly to the image-based model, achieving 58.3% identification accuracy. However, in Lycaenidae, BLAST outperformed images, achieving 91.7% accuracy, and in Coccinellidae, it achieved similar results to the DNA-based DL model, with a success rate of 98.6%.

**Fig. 3.**
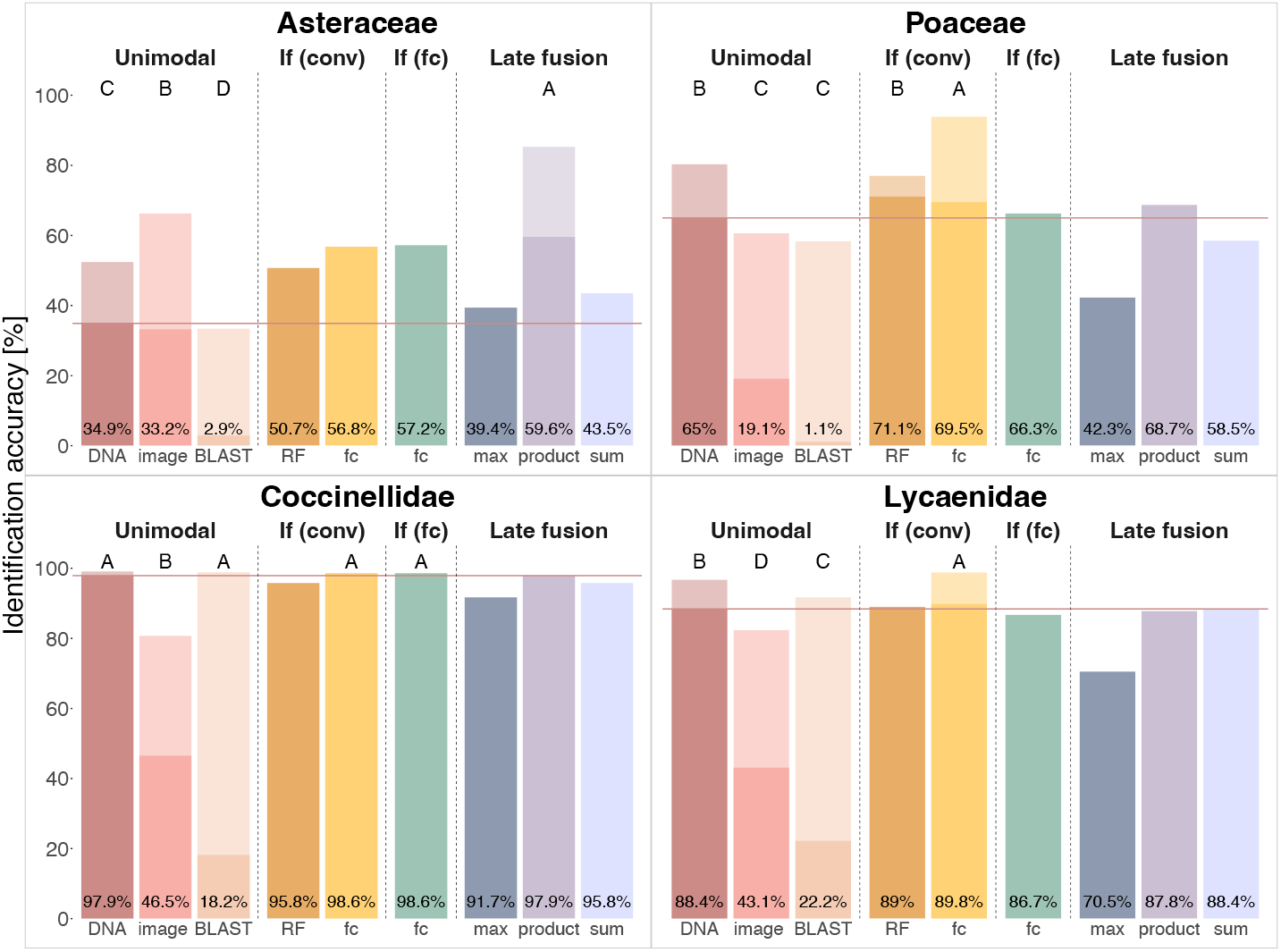
Results of unimodal and multimodal species identification using different fusion approaches: (If (conv), If (fc), Late fusion). Non-LOOCV training results and results of the traditional querying against a BLAST database are shown in saturated bars. The unimodal models, the traditional querying against a BLAST database, and the best fused model(s) were subjected to Leave-One-Out Cross-Validation (LOOCV; shown in light-colored bars). The distribution of identification success was statistically compared, resulting in letters indicating significant differences (paired Cochran’s Q and pairwise McNemar’s tests, p<0.001), where A indicates the best performance and C/D the worst. The solid horizontal line illustrates the identification accuracy achieved by the superior unimodal model during the non-LOOCV training, reinforcing which of the multimodal models outperformed the unimodal models. If intermediate fusion, RF=Random Forest, fc=fully connected.

### Multimodal species identification

Reusing the weights gained from training on a single modality (see Section Training), we combined the two modalities in a second training. In general, the fusion of DNA and image data outperformed unimodal species identification models across all datasets using the initial training and validation split (Fig. 3), with the increase in accuracy being more pronounced for the plant datasets than for the animal datasets. Product score fusion, as well as the intermediate (conv) and intermediate (fc) fusion approaches employing fully connected layers, consistently performed the best across datasets.

We validated our findings by applying LOOCV on both image- and DNA-only models as well as the fusion model(s) that yielded the highest success rate, i.e., product-score for Asteraceae, intermediate (conv) using a fully connected classifier for Poaceae, Coccinellidae, and Lycaenidae, and intermediate (fc) for Coccinellidae. In the Asteraceae dataset, both the DNA-based and image-based models exhibited a lower identification accuracy compared to the multimodal model. The score-level fusion approach using the score product outperformed the best unimodal model, the image-based model, by approximately 19% (p 0.0001). Based on the initial training with the traditional training-validation split, we observed that the best fusion approach for all other datasets was the intermediate fusion (conv) approach using two fully connected layers for classification. In Poaceae, the intermediate fusion (conv) achieved a more accurate species identification compared to DNA alone (p 0.0001) with 93.9% compared to 80.3%, respectively. Even when DNA-based models achieved very high identification success in the unimodal approach, as is the case for Lycaenidae (96.7%), we have been able to exceed the accuracy scores by using the intermediate fusion (conv) approach (98.8%, p 0.0001). Only in Coccinellidae did both selected fusion methods not surpass the DNA-based model significantly. Here, the intermediate (conv) and intermediate (fc) fusion methods, the DNA-based model, and the approach using BLAST resulted in comparable identification accuracies.

For both plant datasets, the baselines were still considerably lower than those of the two animal groups. Thus, in plants, the fusion approaches that outperformed the unimodal models do so by a large margin (Asteraceae.: + 19%; Poaceae: +13.6%). In animals, where DNA-based models achieve very high accuracies on their own, the differences were less pronounced (Lycaenidae: +2.1%).

The expanded LOOCV that included all samples within the Asteraceae and Coccinellidae datasets confirmed the results of the subset-based approach (Fig. S2).

### Inter- and intrageneric confusion

To understand which characteristics of the dataset may lead to low baseline accuracies and to understand what effect the two modalities have on the model performance, we compared intra- and intergeneric confusion rates. We observed middle to low levels of misidentification per species across datasets and models with means ranging from 0% to 37% per dataset (Table 2). In general, plants showed higher levels of confusion compared to animals (p 0.0001). DNA-based and fused models were significantly less prone to confusing samples between genera than image-based models (Asteraceae: p 0.0001, Poaceae: p 0.0001, Coccinellidae: p 0.05, Lycaenidae: p 0.0001). In plants, barcodes and fused data were predominantly confused within rather than between genera (p 0.0001), while images tend to be confused more often between than within genera (p 0.0001). In animals, DNA is rarely confused with only 2.8% of species showing any level of misidentification. Images, however, display the pattern observed in plants and are more frequently misclassified on the level of genus than of species (p 0.05). Across datasets and confusion levels, the fusion approach that yielded the highest identification accuracy delivers comparable results to the unimodal approach that performs better (see Figure S7 for a per-genus perspective on confusion rates). The only exception poses intrageneric confusion of Asteraceae images, where images alone are confused less often than the combined modalities. However, in all other cases, fusion is a combination of the superior result of each modality. When investigating further and looking into duplicate sequences within all four datasets, DNA confusion was shown to be strongly linked to duplicates within and in between genera as revealed by Figure 4 (sample duplicate level: ‘intrageneric’/’intergeneric’). Particularly in Asteraceae and Poaceae, DNA samples were mainly falsely assigned to species that contained a duplicate to the respective sample in the training set (sample duplicate level: ‘combi’). In all datasets, they were often confused with species that exclusively included duplicates and, therefore, had a genetic distance of 0, or with species with very little genetic difference to the DNA sample. Oftentimes, these confusions could be solved by integrating image information since the genetic difference between true and assigned species was much larger when identifying using images. Information on samples that were only correctly identified by a single modality or not correctly identified at all is shown in Figures S8-S10.

**Table 2.** Results of paired Kruskal-Wallis and pairwise Wilcoxon signed rank tests for intergeneric (above) and intrageneric (below) confusion rates between tested modalities. Tests were applied to the confusion found during Leave-One-Out Cross-Validation (LOOCV). Letters indicate significant differences between modalities (A=highest confusion).

|  | Asteraceae | Poaceae | Coccinellidae | Lycaenidae |
| --- | --- | --- | --- | --- |
| **intergeneric confusion** |  |  |  |  |
| Kruskal | 108.67 | 176.9 | 42.1 | 168 |
| df | 2 | 2 | 2 | 2 |
| p-value | <0.0001 | <0.0001 | <0.0001 | <0.0001 |
| modality | mean confusion<sub>LOOCV</sub> |  |  |  |
| barcode | 10.8%<sup>B</sup> | **4.3**<sup>B</sup> | **0.7**<sup>B</sup> | **0.5**<sup>B</sup> |
| image | 25% | 36.8% | 5.9% | 11.9% |
| both | **2.2**<sup>C</sup> | **2.6**<sup>B</sup> | **1.7**<sup>B</sup> | **0.5**<sup>B</sup> |
| **intrageneric confusion** |  |  |  |  |
| Kruskal | 35.09 | 23.01 | 21.78 | 38.73 |
| df | 2 | 2 | 2 | 2 |
| p-value | <0.0001 | <0.0001 | <0.0001 | <0.0001 |
| modality | mean confusion<sub>LOOCV</sub> |  |  |  |
| barcode | 36.8% | 15.4% | **0.3**<sup>B</sup> | 2.8%<sup>B</sup> |
| image | **8.7**<sup>C</sup> | **2.6**<sup>B</sup> | 5.9% | 5.7% |
| both | 12.5%<sup>B</sup> | **3.5**<sup>B</sup> | **0**<sup>B</sup> | **0.7**<sup>C</sup> |

**Fig. 4.**
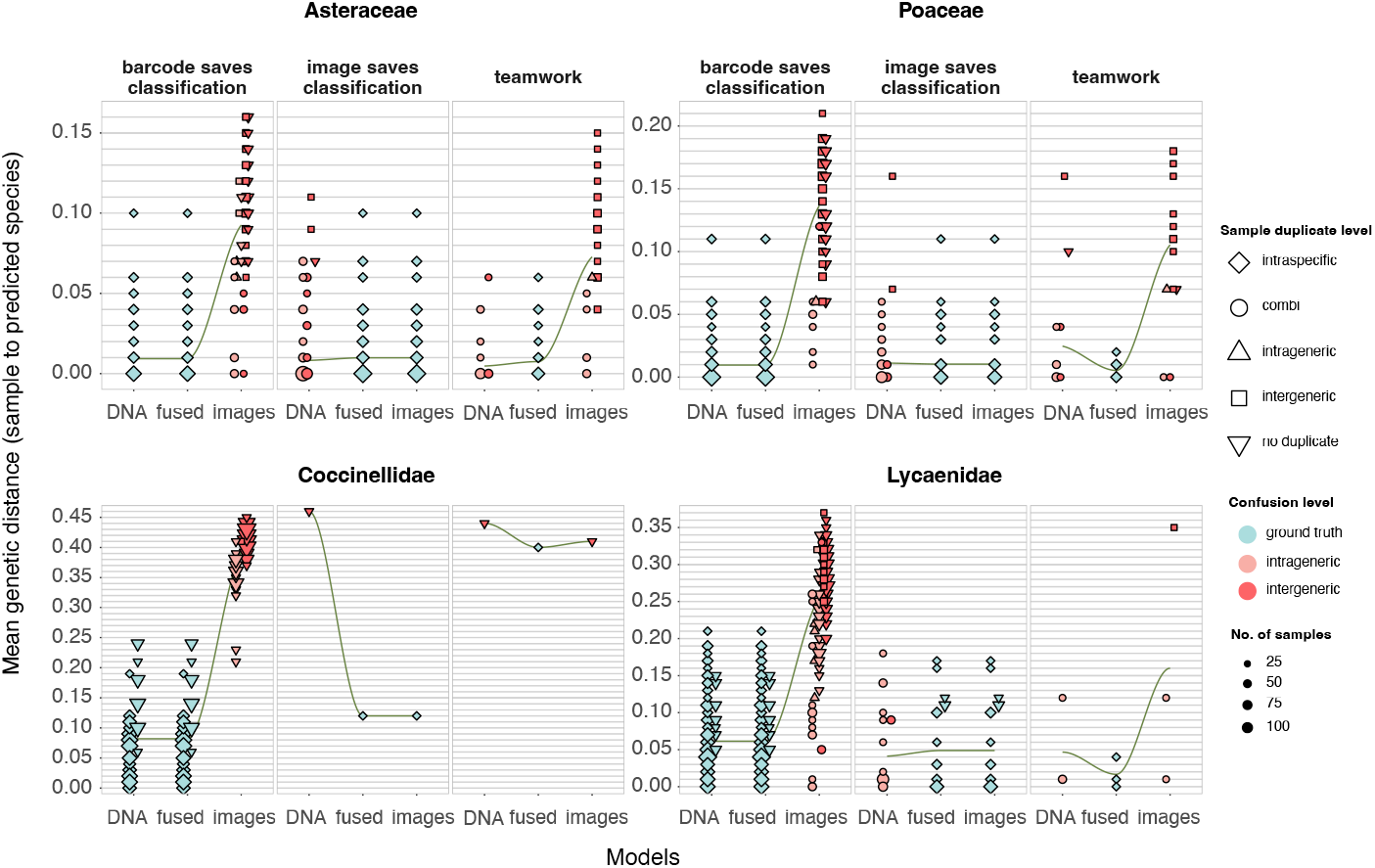
Shown are confused samples that were either correctly identified by a) the barcode-only and (best) fused model, b) only the fused model, or c) the image-only and fused model. The y-axis displays the mean genetic distance (rounded to two decimal places) between the tested sample and the training samples of the predicted species. The level at which the samples were confused is indicated by the color; the shape provides information on whether and where there is a duplicate sequence in the training dataset. Numbers of samples matching genetic distance, duplicate status and confusion level are shown via data point size.

#### Confusion in relation to gene length and sample size

We found significant relationships between inter- and intrageneric confusion rates and number of training samples and species within the genus as well as gene length (Table S1). In detail, fewer samples were misidentified when the number of training samples was larger in the following cases: a) Asteraceae, DNA-based model, intra- and intergeneric (p 0.05), b) Poaceae, image-based model, intrageneric (p 0.05), and c) Lycaenidae, image-based model, inter- and intrageneric (p 0.01). The mean gene length and the number of species within the genus had a significant impact on the confusion rate when examining pooled data from all datasets. Here,, the influence gene length has on confusion was not conclusive in terms of positive/negative impact. For example, while a longer validation gene length increases intrageneric confusion (p 0.001), it decreases intergeneric confusion in DNA-based species identification (p 0.0001). In multimodal species identification, the effect was positive for both intra- and intergeneric confusion (p 0.0001 and p 0.05, respectively). The number of species within the genus increases intrageneric confusion levels of both DNA-based and multimodal models (p 0.001, p 0.01). Lastly, the difference between the gene length in the training set compared to the validation set impacted intrageneric confusion rates for DNA-based and multimodal models. However, while the effect was positive in the DNA-based model (p 0.0001), it was negative in multimodal training (p 0.01).

## Discussion

This study, for the first time, systematically analyzed various DNA preprocessing methods and multimodal fusion approaches. We demonstrated that (i) fusion widely outperforms unimodal identification, with fractional encoding of DNA combined with intermediate (conv) and intermediate (fc) data fusion achieving the highest identification accuracy in three out of four eukaryotic species groups, (ii) fusion significantly improved identification even when genetic data yielded high species identification accuracy, (iii) fusion reduces both high intrageneric confusion of barcode-based identification and high intergeneric confusion of image-based identification.

### DNA preprocessing

To date, there has not been an investigation on the effect of different DNA preprocessing techniques. Yet, DNA preprocessing is a crucial step to ensure the efficiency and effectiveness of a model. In this study, we compared six preprocessing methods. We observed the most accurate species identification when first aligning the sequences before applying fractional encoding in one plant and one animal dataset. This approach is consistent with the practice of the majority of studies dealing with genetic data in the context of ML while contrasting projects that rely on unaligned sequences for analyses (e.g. Zhang et al., 2008; Fiannaca et al., 2018; Yang et al., 2022). Notably, we also found that in Poaceae and Coccinellidae an additional step that reduces the aligned sequences to their SNPs yielded results that were on par with the performance of complete sequences. We conclude that the relative number of SNPs, i.e., the retained information, and its balance with the loss of information that may arise by removing conservative regions is the major factor contributing to the difference in performance between SNPs vs complete alignments in some datasets. In detail, discarding conservative positions can disrupt meaningful patterns that then form simpler patterns that, without the respective context, are much more prevalent across locations within the sequence and between samples. However, the success with using SNPs in Poaceae and Coccellindae shows that SNPs can be as informative as the complete sequence. This finding can contribute to model runtime reduction e orts when dealing with large multi-gene datasets in future research. In addition, the use of a non-CNN architecture that does not rely on recognizing patterns within immediate local surroundings could improve the identification using SNPs. For instance, transformers are a viable option for capturing non-local interactions as they use, in contrast to ResNets, an attention mechanism (e.g. Ji et al., 2021).

#### Uni- and multimodal training

We have provided a fundamental comparison of fusion stages and classifiers that can serve as a basis for future studies seeking a more holistic perspective on species identity when training ML models. With respect to unimodal models, barcode-based models usually yielded higher identification accuracy than image-based models. Notably, the identification of plant species has proven to be much more challenging than the identification of animals, with substantial differences in the ability to classify on barcodes alone. In Asteraceae, the models based on barcodes were significantly outperformed by images. We attribute this to the frequent occurrence of duplicate sequences in our plant datasets, particularly in the Asteraceae dataset, which not only a ects within-species confusion but also misidentification between species or even genera. Events like apomixis, hybridization, and polyploidy may contribute to this circumstance (Fazekas et al., 2009; Karbstein et al., 2024). We discovered, however, that some of the cases in which species are confused due to one or more duplicate sequences in the training set can be resolved by including the information provided by the image. Our results show that the fusion of morphology and genetics is usually beneficial, even when the genetic information itself is sufficient to identify a vast majority of test samples. Fusing genetics with image data significantly outperformed unimodal models for three out of four datasets. In Lycaenidae, fusion after feature extraction with two shared fully connected layers outperformed the barcode-only model by 2% while barcodes alone already classified 97% of samples correctly. While identification accuracy did not increase with fusion in Coccinellidae, it is worth noting that the dataset included fewer species and, at the median, more samples per species compared to all other datasets, potentially rendering the task less difficult for the model. Overall, integrating genetic and image features using a fully connected classifier consistently produced the best or near-best results and therefore be recommended for integrative species identification e orts. An explanation for the improvement brought by fusing genetics with image data is the limited resolution of barcodes that has been discussed several times in the past (Besse et al., 2021; Ahmed et al., 2022). The genes that we used in this study were, on average. 550-650 bp long, while the median lC-value (DNA in a haploid nucleus) of, e.g., angiosperms, is 2.4 Gbp. Furthermore, plant genomes comprise

40,000 genes on average (Sterck et al., 2007). Considering this, a single gene is only a tiny snapshot of the entire genome. In addition, natural selection acts on these DNA fragments, reducing their variability and, in turn, their ability to differentiate closely-related species in particular. In combination with the aforementioned intricate evolutionary processes, these effects may result in large amounts of completely indistinguishable samples (Zarrei et al., 2015; Karbstein et al., 2022). The stark contrast between accuracies achieved by animal versus plant DNA-based models can be attributed to the specific markers used in this study. When barcoding animals, *COI* represents the consensus due to its discriminatory power (Hebert et al., 2003b; Ahmed et al., 2022). However, in plants, the mitochondrial gene *COI* shows lower variation because it evolves too slowly in plants, therefore, nuclear and plastid genes are used more often (Hollingsworth et al., 2011). Furthermore, it has been shown that one marker alone tends to not be sufficient to distinguish between species (Hollingsworth et al., 2016). Two or three markers are commonly used in conjunction to provide fine-grained resolution (e.g. Romeiro-Brito et al., 2016). Consequently, a substantially higher identification accuracy in animals compared to plants when using only one plant marker is to be expected. Notably, the potential of an integrative approach to species identification depends on the information already Contained within each modality. Researchers should choose carefully when opting for a multimodal, more time and resource consuming approach by first assessing the relative gain of such a method, particularly when working with animal DNA. Yet, the remaining, not sequenced DNA the network is not trained on as well as environmental factors that reflect in epigenetics can be discernible by the network through a condensed manifestation in the morphology of the specimen. Our findings reinforce the direction proposed by Karbstein et al. (2024) for integrative species delimitation and that there is a need for, at least, utilizing a multitude of genetic and morphological information as well as metadata for accurate species delimitation (i.e., species delimitation 3.0). The identification accuracy achieved by using herbarium material for the Asteraceae dataset proves that specimens from collections are a valuable data source for integrative taxonomic; ML approaches, Collection data has already started to gain traction in biological ML research. For example, studies focusing on phenological stage identification (Pearson et al, 2020; Katal et al., 2022), and plant organ segmentation (Weaver and Smith, 2023) leverage herbarium material. Given that museum samples are reliably labeled, even supervised learning algorithms can be applied without further work necessary. Recently, features learned by an ML network and geometric morphometrics-based features extracted manually from both *in situ* and herbarium specimen images have been shown to be significantly correlated (Hodač et al., 2024), demonstrating that ML is able to learn meaningful features from herbarium samples. Use of collection material allows for cheaper studies with larger datasets and, potentially, more robust results and should therefore be considered when working with ML. An important aspect of this study is the usage of independent data points for DNA and images. Studies such as Yang et al. (2022) use co-occurring data, i.e., DNA and image originate from the same individual. Dependent data ensures that the variance experienced by the model is part of the naturally occurring distribution, which may lead to better generalization when confronted with other samples of the; same distribution. Yet, this poses a vital problem as co-occurring data can be hard to come by and, thus, can further limit and complicate expensive dataset collection e orts. Consequently, no sampling e ort will ever cover all variance encountered within the naturally occurring distribution. When working with images, a common procedure is data augmentation to semi-artificially increase the dataset size and introduce more variation, which leads to better generalisation. Even when working with DNA, data augmentation can improve model performance (Lee et al., 2023). When working with multiple modalities, a way to augment the data can be to shuffle the modalities independently, thereby creating more variation. Co-occurring data should therefore not be a hard requirement for sampling e orts.

#### Confusion

Confusion patterns differ between images and barcodes. Images are oftentimes confused between genera while barcodes tend to be confused within genera. In addition, our findings suggest that datasets that contain species with significant genetic overlaps, i.e., in cases where the barcoding gap is nearly or completely nonexistent, benefit the most from inclusion of additional modalities. In those cases, these confusions could be solved by integrating image information, highlighting the usefulness of an integrative taxonomic approach to machine learning (Derkarabetian et al., 2019; Alexander Pyron, 2023;: Karbstein et al., 2024). Furthermore, cases in which either or both the molecular data and the image alone did not suffice for a correct prediction but succeeded when used in tandem hint towards a hierarchical role of the molecular data in the identification process. The barcode may guide the model to the correct genus and then settle on the correct species with the help of the image.

#### Limitations

The choice and quality of the genetic markers is an essential prerequisite to the success of fusion approaches using ML. As seen, fusion was not able to outperform the barcode-based approach in Coccinellidae as the baseline resolution provided by the barcodes was close to perfect for the species in our dataset. The barcodes used in this study were chosen based on the number of samples found in freely available online repositories. We did not assess multiple barcodes for, e.g., the plant datasets, to confirm that no other genetic marker would be better suited for species discrimination. However, *rbcLa* is widely used in plant research, oftentimes in combination with *matK* (Li et al., 2015). In addition, taxonomically challenging groups such as those where apomixis, hybridization, and/or ploidy are prevalent pose a significant challenge to all plant barcodes (Fazekas et aL,. 2009; Hollingsworth et al., 2016). We have shown that fusion can improve discrimination even in groups where duplicate sequences are common. Therefore, irrespective of the marker used in this study, we believe that these findings can be universally applied where species are hard to distinguish by genetics alone. DL algorithms typically require substantial data to perform effectively. While we acknowledge that the datasets our models were trained on are relatively small, given the scarcity of many species groups, it becomes imperative to explore and understand how ML models operate under conditions of limited data availability. We are aware that the sample sizes of our datasets are not ideal for training DL models, but believe that working with a limited amount of data is essential, as it reflects the reality of taxonomy-focused studies (Karbstein et al., 2020; Klasen et al., 2022; Opatova et al., 2024). Furthermore, our focus was not on the absolute accuracies attained by our networks, but rather on the relative gains and losses between uni- and multimodal models and different fusion strategies. Given that all models had access to the same number of records, we consider the relative results to be unaffected by the total sample size. Imbalanced sample sizes across classes, as is the case with all our datasets, cause some features/classes to be trained more often than others, leading to model bias. However, as all tested models were trained on the same data, model performance is comparable. We also sampled our validation set evenly across classes to get a balanced look at identification success. In addition, the learning process of ML models is still a ‘black box’ for human observers, Although new explainable Al approaches are emerging to visualize and detect biological features learned by the ML model (Samek et al., 2021; Hodač et al., 2024), previous delimitation results from integrative taxonomy are needed, as well as experienced taxonomists to control and validate ML-based species identification or clustering.

## Conclusions

Modern integrative taxonomy aims to combine 21st century high-throughput sequencing (genomics) methods with multiple complementary data sources, such as morphology from geometric morphometries, ploidy reproduction from flow cytometry, physiology from biochemical screenings, behavior from camera field observations, or biogeography/ecology from environmental statistical modeling (Dayrat, 2005; Schlick-Steiner et al., 2010; Karbstein et al., 2024). Modality-specific shortcomings are reduced in this way, for example single, few, or even hundreds of genes are often not variable enough to differentiate closely-related species (e.g. Tomasello et al., 2020; Dietz et al., 2023) and therefore images of field or herbarium specimens, ploidy, reproductive, behavioral, or ecological niche information can help to add subtle features for more accurate and reliable species identification. As integrative taxonomy is a major avenue for meeting the nature of species in (semi-)manual species delimitation (Dayrat, 2005; Schlick-Steiner et al., 2010; Karbstein et al., 2020, 2022), the joint use of modalities should also be considered a crucial pillar of any ML-based or -assisted approach to identification. The rationale behind this lies in the fact that the taxonomic labels of the underlying dataset are derived from the evaluation of multiple modalities. Consequently, future ML studies should focus on evaluating >2 datasets to test generalizability across taxonomic groups and >1 modality, preferably including >1 genetic marker to reflect delimitation procedures.

This study paves the way by demonstrating, for the first time, that DNA+image fusion strategies merging the features directly after the last convolution tend to yield the best species identification success-Modern integrative taxonomic approaches produce many, and often extraordinarily large datasets, which regularly cannot be handled by traditional phylogenetic and statistical methods in time-efficient ways. DL approaches have the advantage of automatically extracting and concentrating the most important, even complex or subtle features not visible to the human eye for identification from extremely large data matrices in short time frames (Borowiec et al., 2022; Badirli et al., 2023). Consequently, future developments in data fusion are likely to accelerate integrative taxonomic workflows for species identification and delimitation.

## Data Availability

The authors declare that basic data supporting the findings will be available upon publication.

