## Appendix for "Data Fusion for Integrative Species Identification Using Deep Learning"

#### *Text S1: Random Forest Grid Search*

We chose Random Forest parameters by applying RandomizedSearchCV included in the model selection of scikit-learn (Pedregosa et al., 2011). In total, we tested the number of estimators (100-3000), the maximum number of features per leaf (log2 and sqrt), the maximum depth of the trees (10-110), minimum samples per split (2, 5, 10, 15) and per leaf (1, 2, 4), and bootstrapping (True, False). The optimal parameters for our dataset were 1046 estimators, at least 2 samples per split, at least 1 sample per leaf, maximum features defined by log2, a maximum depth of 63, and no bootstrapping.

11 3

*Species identi cation stability*

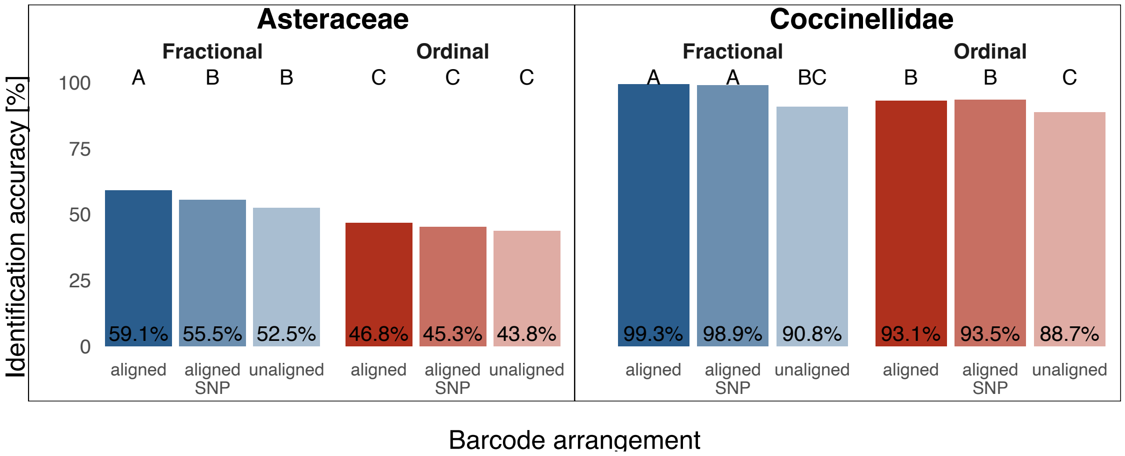


Fig. S1. DNA-based species identification accuracy after di erent arrangement (aligned, aligned-SNP, unaligned) and encoding (fractional, ordinal) methods for genetic data for all samples within the two datasets Asteraceae and Coccinellidae. Identification accuracy describes the percentage of samples within the validation set correctly identified by the model. Letters indicate significant di erences in performance (paired Cochran's Q and pairwise McNemar's tests, p<0.001). SNP = Single Nucleotide Polymorphisms.

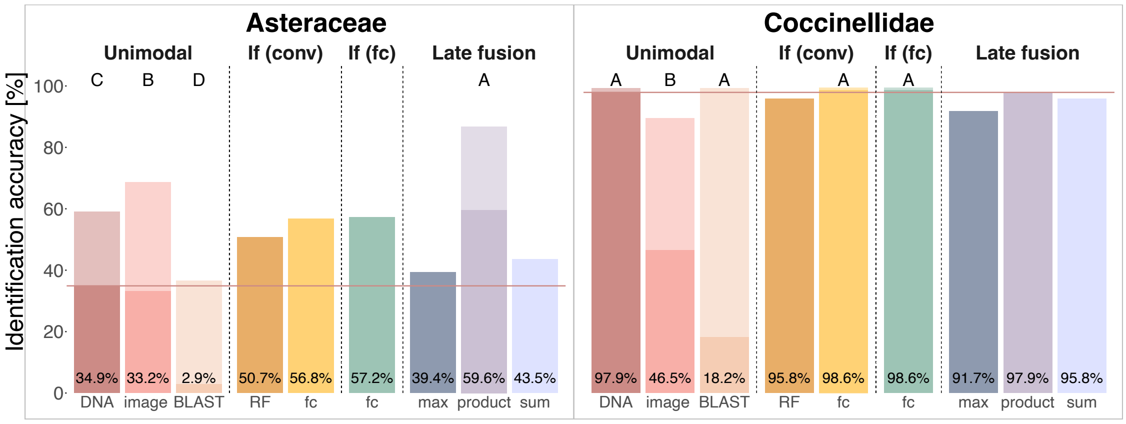


Fig. S2. Results of unimodal and multimodal species identification using di erent fusion approaches (If (conv), If (fc), Late fusion). Non-LOOCV training results and results of the traditional querying against a BLAST database are shown in saturated bars. The unimodal models, the traditional querying against a BLAST database, and the best fused model(s) were subjected to Leave-One-Out Cross-Validation (LOOCV; shown in light-colored bars) using all samples within the two datasets Asteraceae and Coccinellidae. The distribution of identification success was statistically compared, resulting in letters indicating significant di erences (paired Cochran's Q and pairwise McNemar's tests, p<0.001), where A indicates the best performance and C/D the worst. The solid horizontal line illustrates the identification accuracy achieved by the superior unimodal model during the non-LOOCV training, reinforcing which of the multimodal models outperformed the unimodal models. If=Intermediate fusion, RF=Random Forest, fc=fully connected.

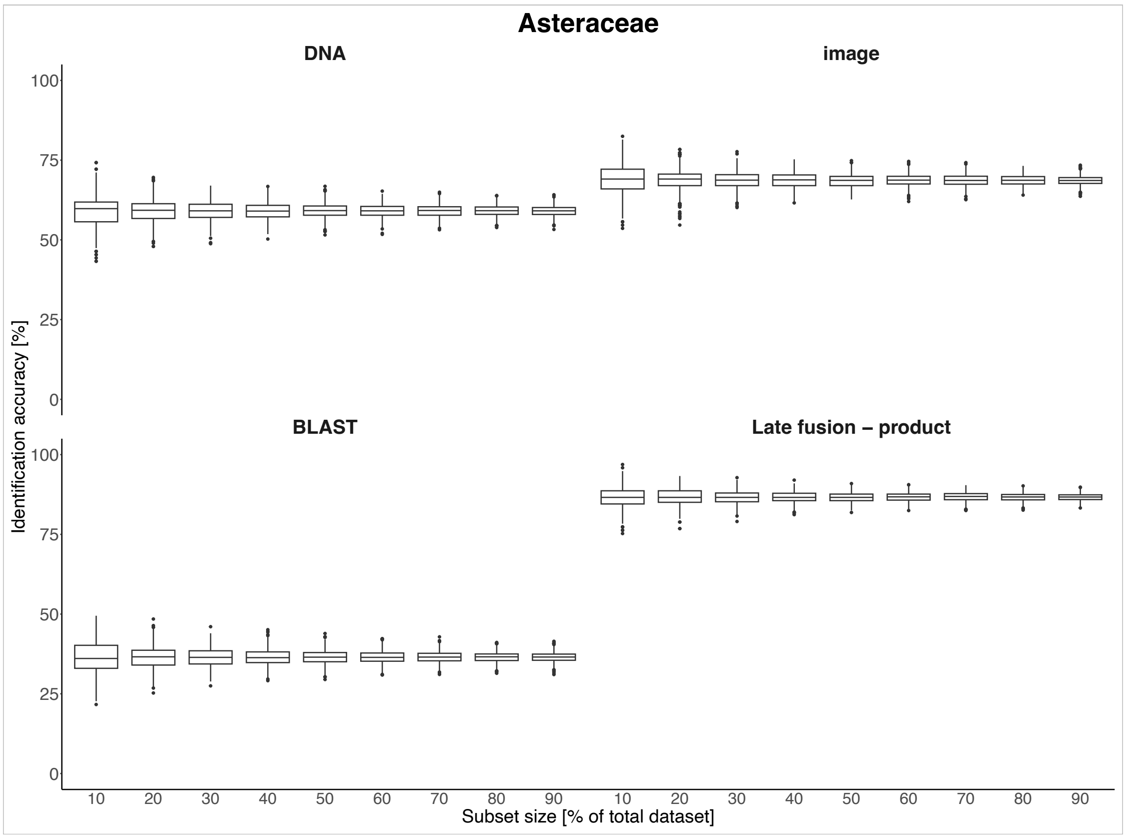


Fig. S3. Shown are the mean accuracies achieved when bootstraping the expanded LOOCV results with replacement based on specific percentage of the entire dataset (10-90% in steps of 10%). We included 1000 repetitions per subsample percentage.

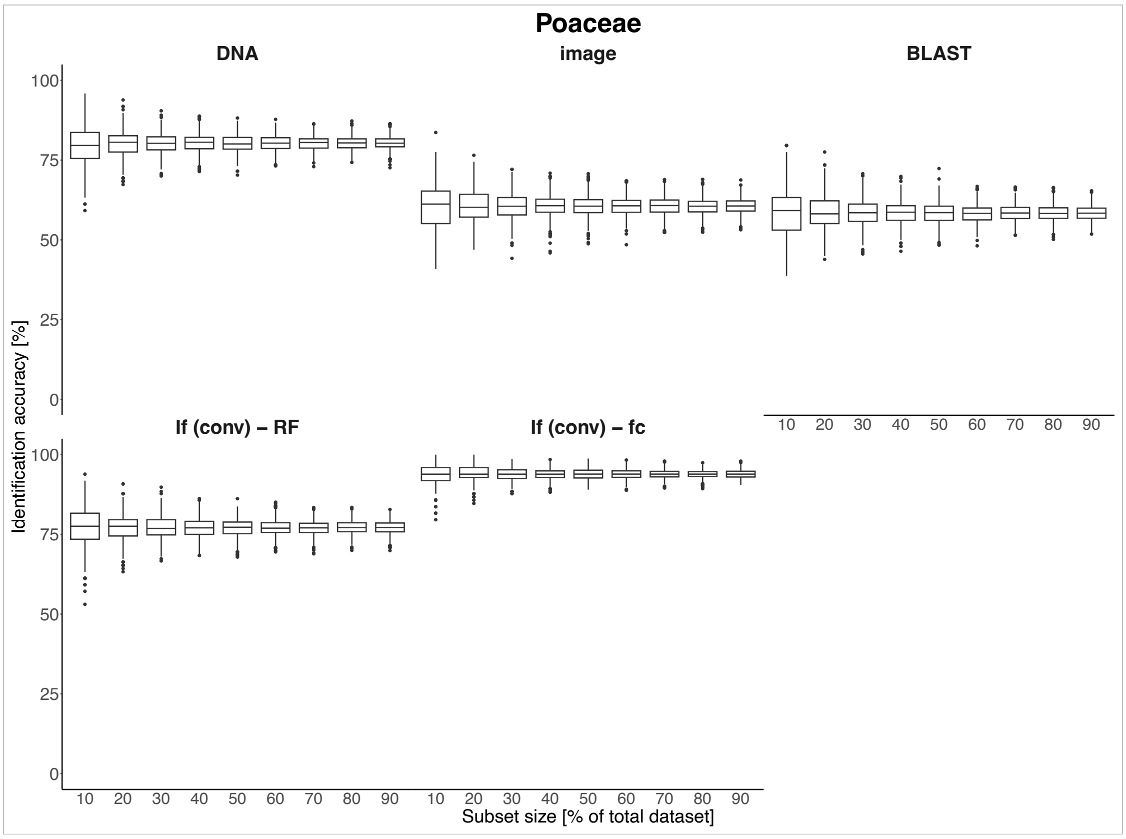


Fig. S4. Shown are the mean accuracies achieved when bootstraping the LOOCV results with replacement based on specific percentage of the entire dataset (10-90% in steps of 10%). We included 1000 repetitions per subsample percentage.

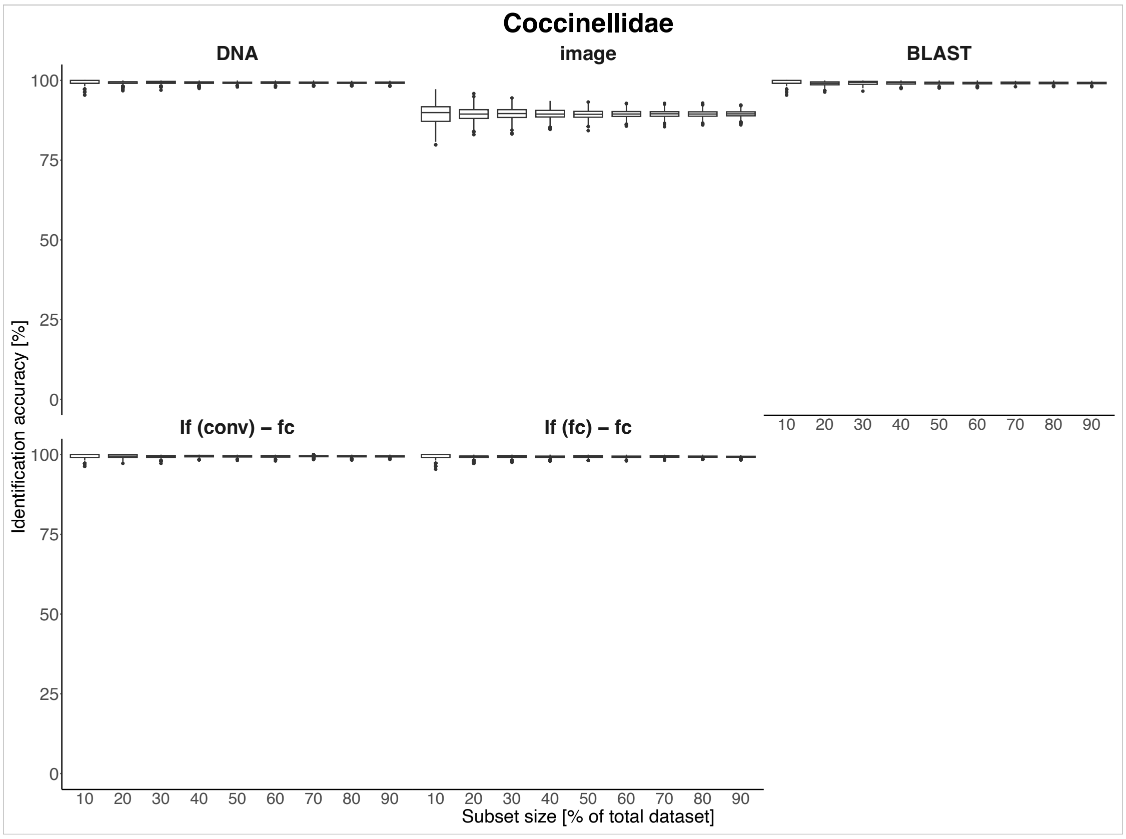


Fig. S5. Shown are the mean accuracies achieved when bootstraping the expanded LOOCV results with replacement based on specific percentage of the entire dataset (10-90% in steps of 10%). We included 1000 repetitions per subsample percentage.

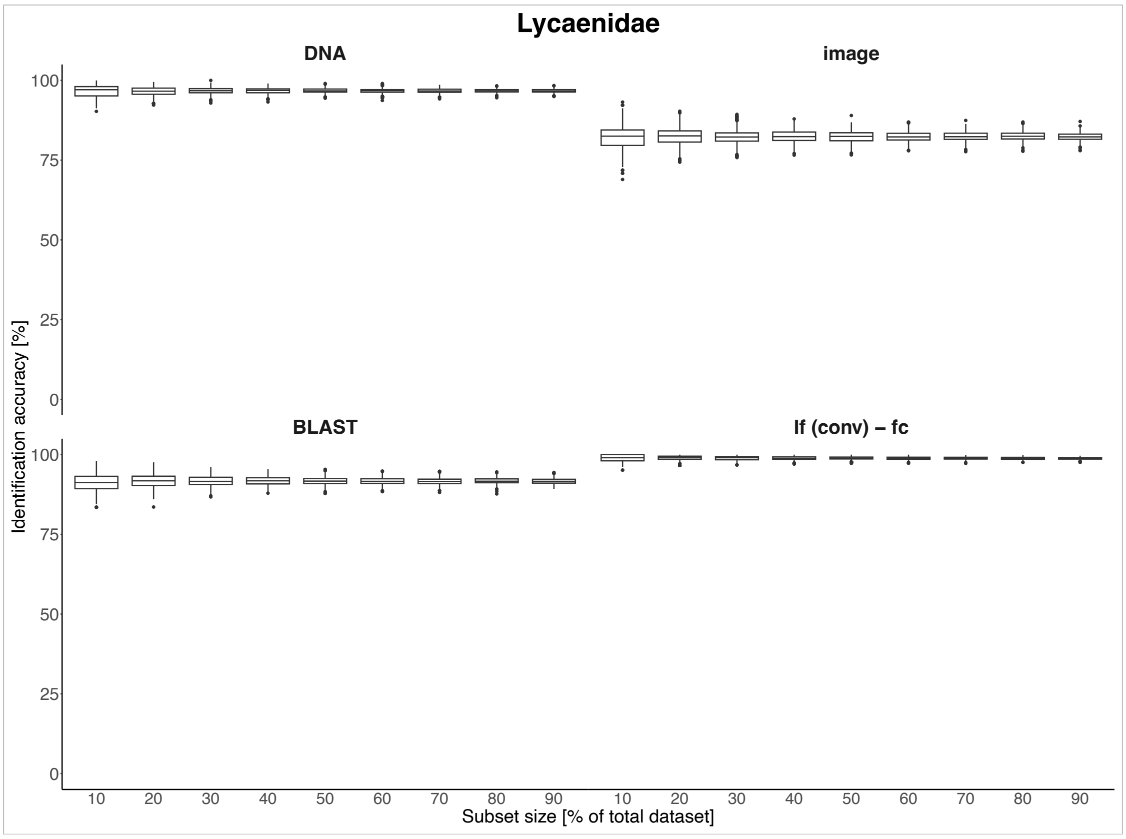


Fig. S6. Shown are the mean accuracies achieved when bootstraping the LOOCV results with replacement based on specific percentage of the entire dataset (10-90% in steps of 10%). We included 1000 repetitions per subsample percentage.

11 4

*Inter- and intrageneric confusion*

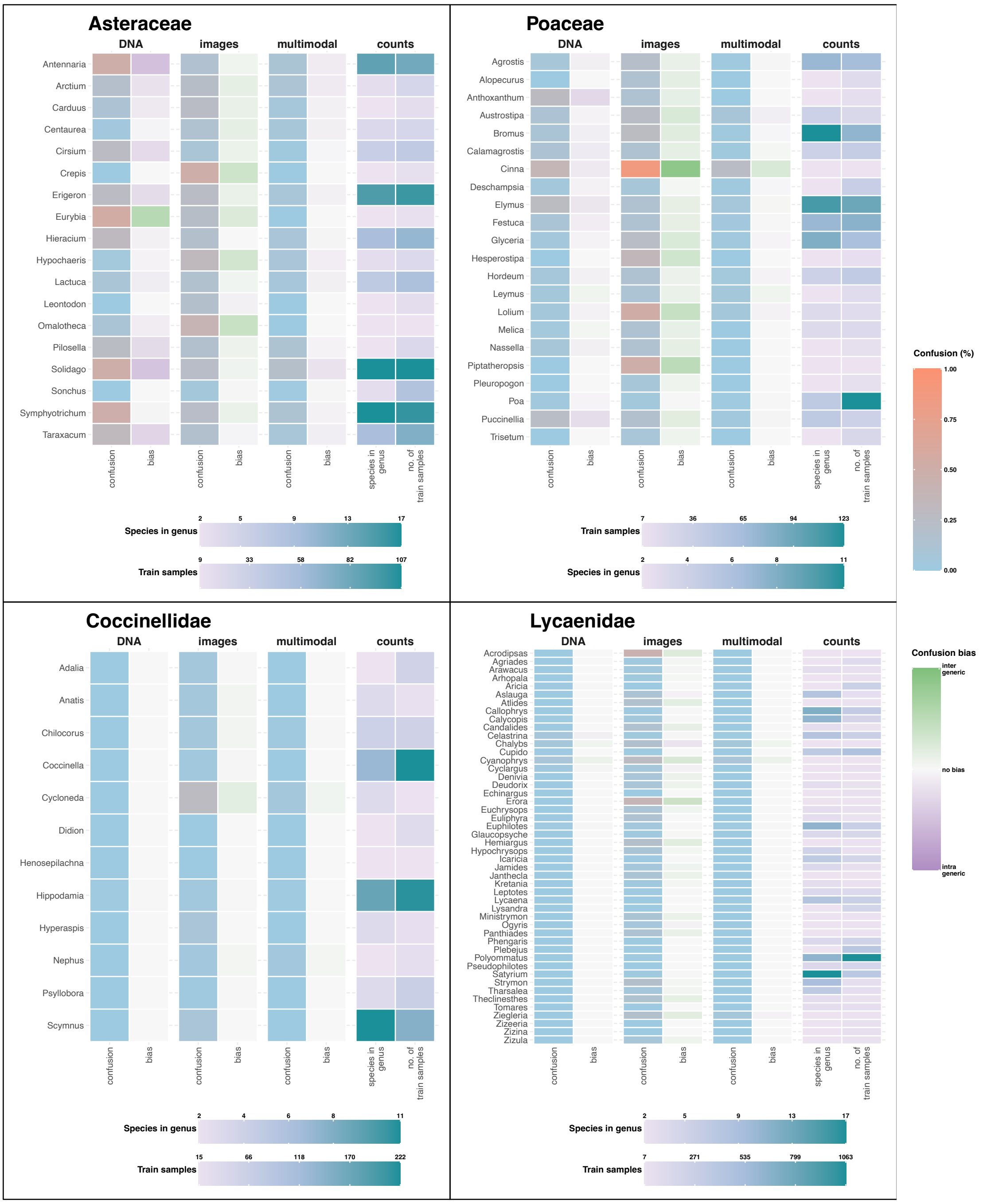


Fig. S7. Shown are confusion rates per genus alongside bias towards inter- or intrageneric confusion for each of the unimodal and the best multimodal models. Additionally, the number of species within the genus and training samples are included.

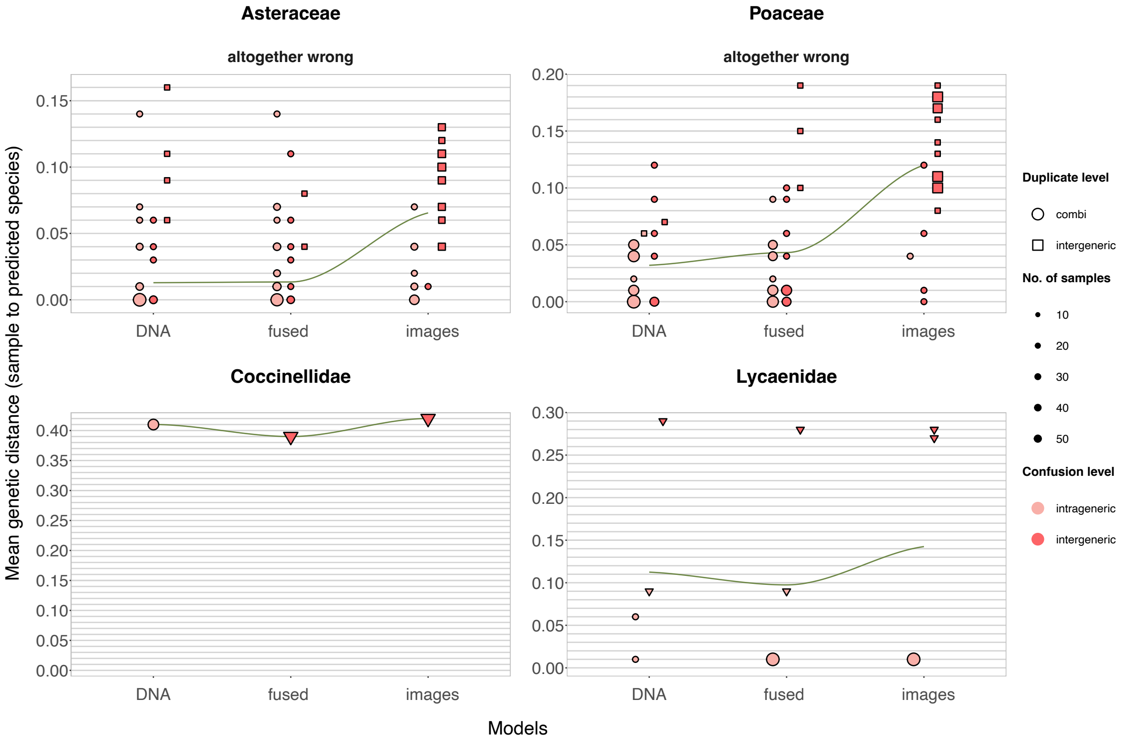


Fig. S8. Shown are confused samples that were misidentified by the barcode-only, image-only, and (best) fused model. The y-axis displays the mean genetic distance (rounded to two decimal places) between the tested sample and the training samples of the predicted species. The level at which the samples were confused is indicated by the color; the shape provides information on whether and where there is a duplicate sequence in the training dataset. Numbers of samples matching genetic distance, duplicate status and confusion level are shown via data point size.

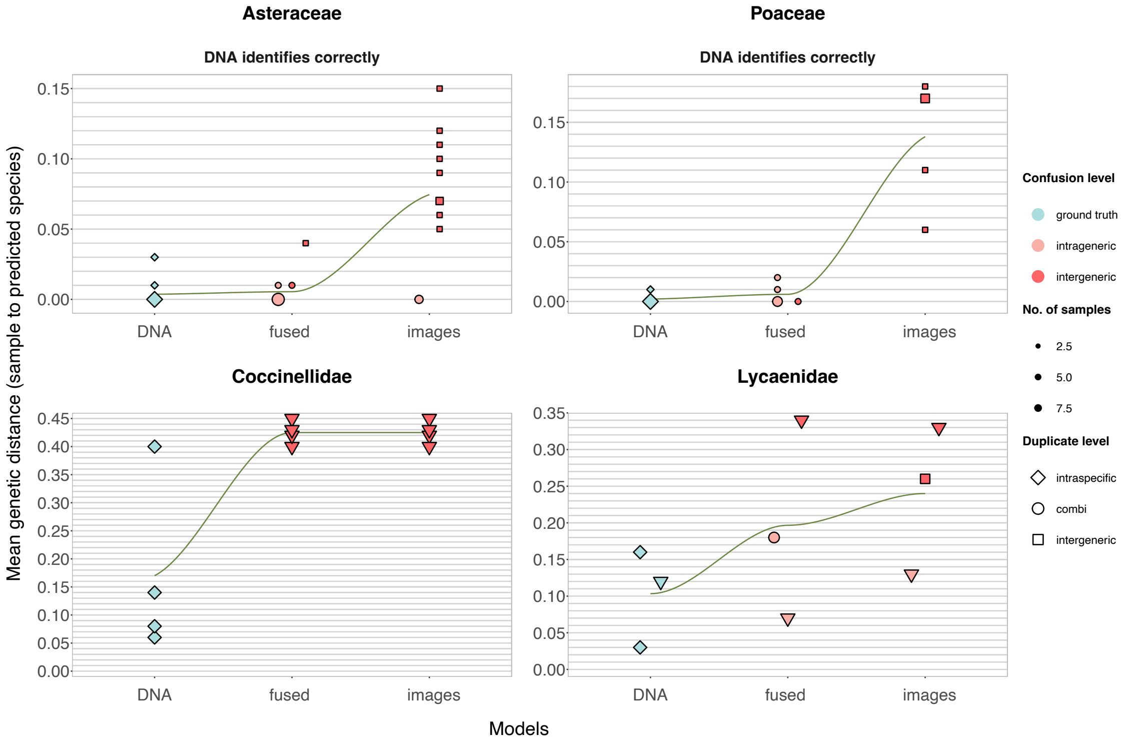


Fig. S9. Shown are confused samples that were misidentified by the barcode-only, image-only, and (best) fused model. The y-axis displays the mean genetic distance (rounded to two decimal places) between the tested sample and the training samples of the predicted species. The level at which the samples were confused is indicated by the color; the shape provides information on whether and where there is a duplicate sequence in the training dataset. Numbers of samples matching genetic distance, duplicate status and confusion level are shown via data point size.

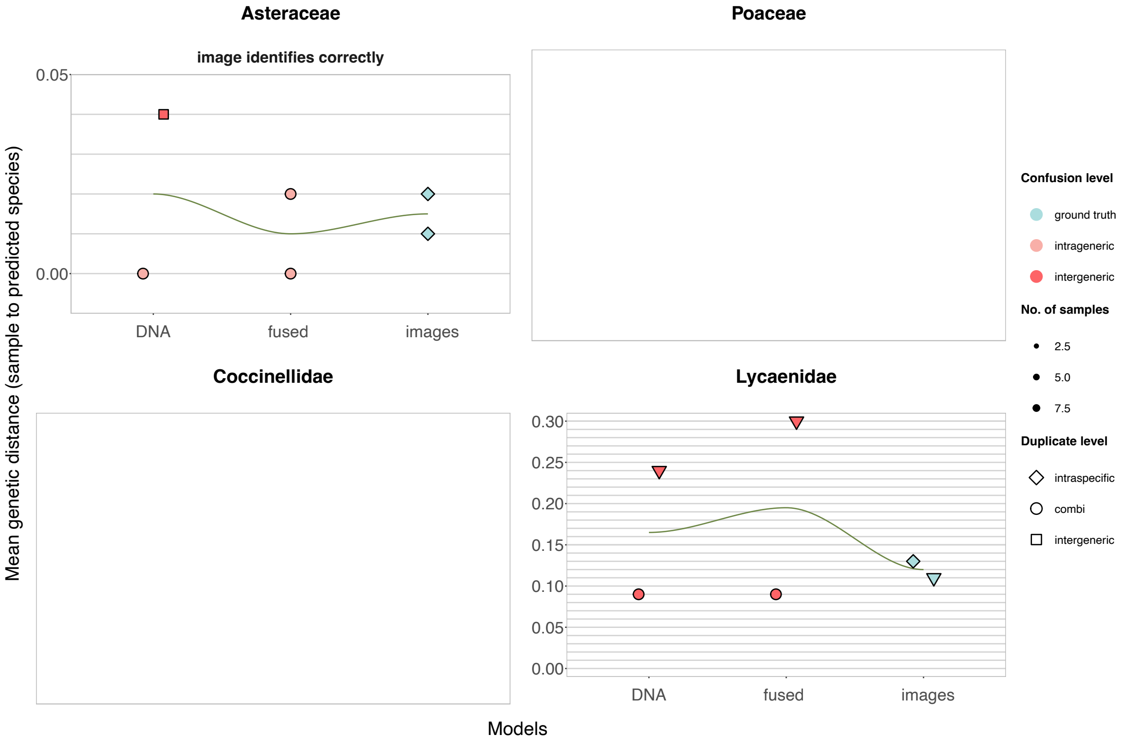


Fig. S10. Shown are confused samples that were misidentified by the barcode-only, image-only, and (best) fused model. The y-axis displays the mean genetic distance (rounded to two decimal places) between the tested sample and the training samples of the predicted species. The level at which the samples were confused is indicated by the color; the shape provides information on whether and where there is a duplicate sequence in the training dataset. Numbers of samples matching genetic distance, duplicate status and confusion level are shown via data point size.

11 5

### *GLM*

Table S1. Response-predictor relationships observed using generalized linear models (GLMs) with binomial response. Statistics were calculated separately for each group, pooled per kingdom, or pooled overall. Train=training dataset, val=validation dataset.

| **group** | **modality** | **response (confusion)** | **predictor** | **estimate** | **p-value** |
|---|---|---|---|---|---|
| Asteraceae | DNA | intergeneric | no. of samples (train) | -0.31555 | <0.05 |
| Asteraceae | DNA | intrageneric | no. of samples (train) | -0.11926 | <0.05 |
| Poaceae | image | intrageneric | no. of samples (train) | -0.06211 | <0.05 |
| Lycaenidae | image | intergeneric | no. of samples (train) | -0.02567 | <0.01 |
| Lycaenidae | image | intrageneric | no. of samples (train) | -0.01789 | <0.01 |
| plants | DNA | intergeneric | no. of samples (train) | -0.13865 | <0.05 |
| plants | DNA | intrageneric | no. of samples (train) | -0.10715 | <0.01 |
| plants | multimodal | intrageneric | no. of samples (train) | -0.09789 | <0.01 |
| animals | image | intergeneric | no. of samples (train) | -0.0237 | <0.01 |
| animals | image | intrageneric | no. of samples (train) | -0.01711 | <0.01 |
| overall | DNA | intergeneric | mean gene length (val) | -0.01045 | <0.001 |
| overall | DNA | intergeneric | no. of samples (train) | 0.14167 | <0.01 |
| overall | DNA | intrageneric | mean gene length (val) | 0.01685 | <0.0001 |
| overall | DNA | intrageneric | species in genus | 0.18419 | <0.001 |
| overall | DNA | intrageneric | no. of samples (train) | -0.03514 | <0.05 |
| overall | DNA | intrageneric | gene length di erence train to val | 0.01202 | <0.0001 |
| overall | image | intergeneric | no. of samples (train) | -0.02518 | <0.001 |
| overall | image | intrageneric | no. of samples (train) | -0.01922 | <0.001 |
| overall | multimodal | intergeneric | no. of samples (train) | -0.19361 | <0.01 |
| overall | multimodal | intergeneric | mean gene length (val) | -0.00704 | <0.05 |
| overall | multimodal | intrageneric | mean gene length (val) | -0.01361 | <0.0001 |
| overall | multimodal | intrageneric | species in genus | 0.13754 | <0.01 |
| overall | multimodal | intrageneric | gene length di erence train to val | -0.01014 | <0.01 |
